# A model for PIP2/3 and Rnd1 effects on Plexin-B1 GAP activity on Rap1b GTPase derived from molecular dynamics simulations

**DOI:** 10.64898/2026.07.09.737506

**Authors:** Nisha Bhattarai, Amita R. Sahoo, Matthias Buck

## Abstract

Plexin-B1 is a transmembrane receptor that integrates signals from Rho-family and Ras-family (Rap1b) GTPases to regulate cellular processes. While ligand simulated activation of the receptor is largely understood, the role of membrane composition and GTPase allosteric effects on plexin structure, internal protein dynamics, and function is still to be elucidated. Here, we performed multi-replica, 1 μs all-atom simulations of Plexin-B1-GTPase complexes on PIP2- and PIP3-containing membranes to investigate the effects of these two signaling lipids, as well as on the GTPases. We found that both Rap1b and Rnd1 stably associate with the membrane, with PIP2 promoting broader lipid engagement and stronger Rap1b-Plexin-B1 interactions, whereas PIP3 enhances Rnd1-Plexin contacts and induces a membrane proximal orientation of Plexin’s juxtamembrane helix and makes contacts with a previously discovered activation switch loop. Contact map and network analyses revealed lipid-dependent shifts in allosteric communication, with PIP2 favoring Rap1b-centric hotspots and PIP3 favoring Rnd1-centric pathways. These predictions allow us to suggest a model for plexin intracellular region activation where both the identity of phosphoinositides and GTPase context synergistically stabilize Plexin-B1 membrane engagement, alter structural dynamics, and allosteric networks. Thus, we propose that the membrane is an active modulator of plexin receptor signaling.

## Introduction

Plexins, which consist of nine family members, serve as receptors for semaphorin ligands and transduce signals essential for neuronal development as well as other processes such as cell adhesion, migration, and immune responses^1–6^. Dysregulation of plexin signaling has been linked to several pathological conditions, including neurological disorders and cancer, with several identified in prostate and breast cancer cells in particular^7–9,19,20^. Structurally, plexins comprise three major regions: an extracellular segment that mediates semaphorin ligand binding, a single transmembrane helix, and an intracellular segment^10^. A unique feature of plexins as single pass-transmembrane proteins (type-I receptors) is their ability to interact directly with small GTPases through two intracellular regions: the Rho-GTPase binding domain (RBD) and GTPase-activating protein (GAP) domain. The plexin GAP domain shares structural similarity with RasGAPs and can interact with Ras-GTPase Rap1b (Fig. 1a)^4,11^. Indeed, one function of plexin involves the hydrolysis of GTP to GDP nucleotide in Ras GTPases^9,12,13^, when these bind the GAP domain as substrates. In addition, plexins interact with several Rho-family GTPases via the Rho-GTPase binding domain (RBD), including Rnd1, Rac1 and RhoD^14,15^. These interactions are thought to play a critical role in regulating plexin activity, with Rnd1 and Rac1 being studied as modulators of plexin signaling^16–18^.

**Figure 1:**
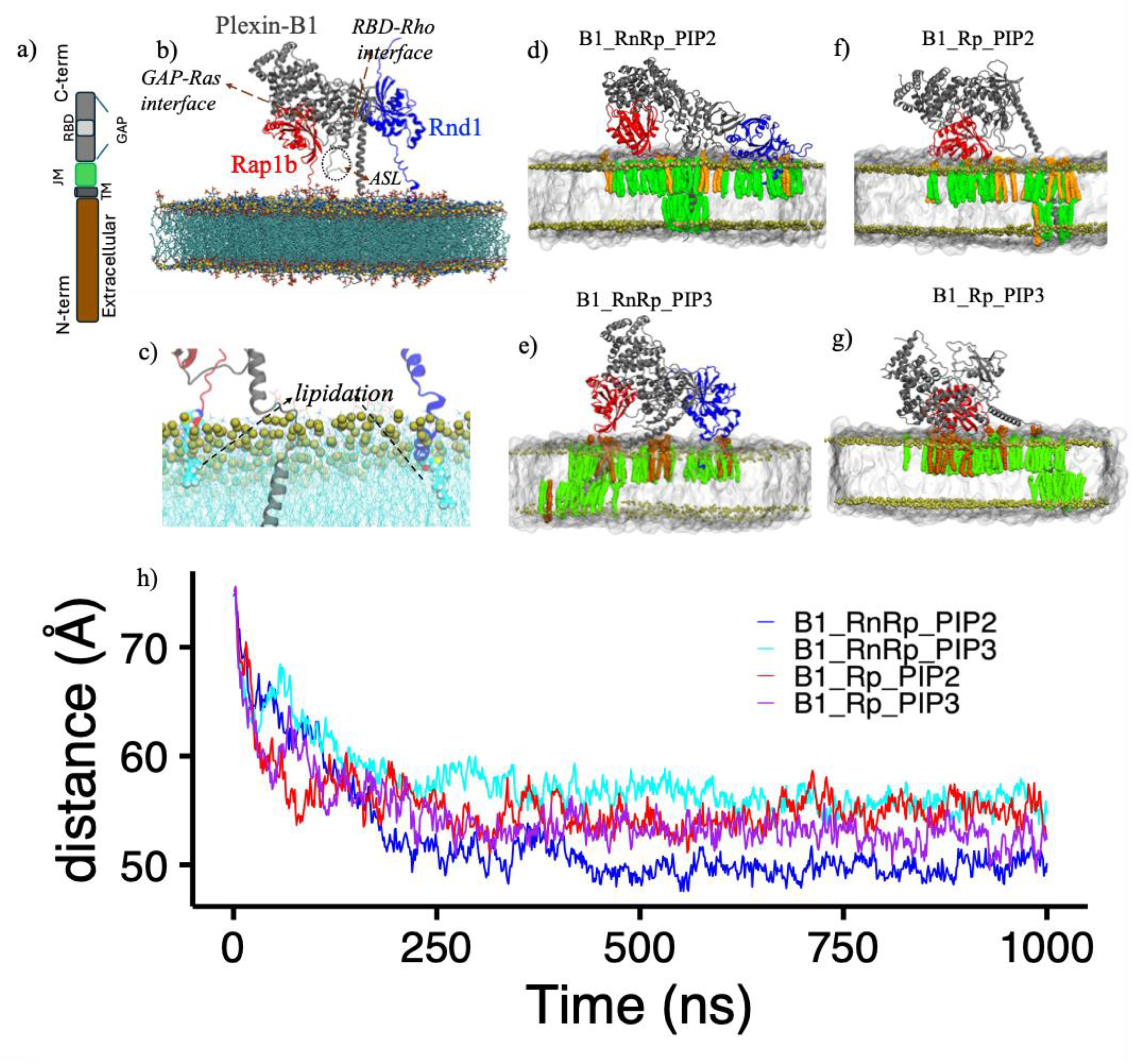
a) Schematic representation of various domains in Plexin-B1. b) Representative initial system setup for Plexin-B1-Rnd1-Rap1b complex, where each protein is colored differently. c) zoom-in view for lipid modification of Rnd1 C-terminal at position C229 with farnesyl modification and Rap1b C-terminal at position C181 with geranylgeranyl modification. d-g) snapshots at the end of ∼1-us simulation where PC/PIP2/PIP3 lipids interacting with the protein are colored green/orange/mustard, respectively, for all complexes. h) Time evolution of the Z-axis center-of-mass distance between the phosphate atoms and the protein center of mass for all systems (Average across three replicas is shown). The average COM distances (mean ± standard deviation across replicas) during the equilibrated portion of the trajectories (last 500 ns) were: B1_RnRp_PIP2: 50.3 ± 1.4 Å, B1_RnRp_PIP3: 56.7 ± 1.6 Å, B1_Rp_PIP2: 55.1 ± 1.6 Å, and B1_Rp_PIP3: 53.5 ± 1.6 Å.

Beyond their canonical nucleoid-dependent conformational and signaling switch, their interactions with regulatory and effector proteins, small GTPases typically associate with membranes, via post-translational lipid modifications, such as farnesylation or geranylgeranylation of their C-terminal cysteines.^19^. Beyond simple membrane attachment, different orientation states are also possible with respect to the bilayer which either occludes or exposes their catalytic/conformational switch domain^20–22^. Thus, it is now appreciated that proper membrane localization and orientation of these GTPases with respect to their binding sites and binding partners is critical in facilitating signal transduction and modulating interactions with effector proteins^23,24^

Despite this, the structural and dynamic determinants governing interactions between lipid-anchored GTPases in complex with plexins and membranes of varying lipid composition is essentially unstudied. Lipid composition of the bilayer, particularly the presence of phosphoinositides such as 4,5-bisphosphate (PIP2)– and phosphatidylinositol 3,4,5-trisphosphate (PIP3), can influence the membrane association of both GTPases and likely Plexin-B1, thereby affecting allosteric communication and functional dynamics within the complex. While this is currently under experimental investigation in the Buck lab, here, we present a computational analysis and predictions from all-atom molecular dynamics simulations.

In our recent work, we studied how the presence or absence of Rap1b at the GAP domain alters the interaction network between plexin-B1 and Rnd1/Rac1 in solution, both experimentally (via hydrogen-deuterium exchange as detected by mass spectrometry, HDX-MS) and computationally^25^. In this work, we investigate how membrane association and lipid composition may modulate plexin-GTPase dynamics at the amino acid residue level. This topic is of importance because, while in cell based experiments there is a clear effect of Rho GTPase Rnd1 binding to the RBD on plexin activity^5^, no crystallography-based studies in the plexin-B1 intracellular domain have been able to delineate a role for these interactions^10^, except for a structure of plexin as a trimer in one crystal form bound to Rnd1 crystallographic trimer ^15^.

To investigate the extent of conformational and dynamic allostery and interactions with lipids, we performed all-atom molecular dynamics simulations of four different systems totaling 12 μs of combined simulation time. Our results show that plexin-B1 and both GTPases (Rnd1 and Rap1b) form more extensive lipid interactions with PIP2-containing membranes than in PIP3-containing membranes, leading to enhanced overall system stability in the former case. Notably, the activation switch loop, a segment between the RBD and the second part of the GAP domain, which was previously identified by computational and experimental studies in our lab ^9^, is central to these interactions. We also find that Rap1b–lipid interactions are modulated by the presence of Rnd1, indicating synergistic effects between RBD- and GAP-bound GTPases. Furthermore, membrane composition (PIP2 versus PIP3) significantly altered residue-level contacts and interaction networks among plexin-B1, Rnd1, and Rap1b, highlighting a previously underappreciated role of the lipid environment in regulating plexin–GTPase signaling dynamics.

## Material and Methods

Construction of Protein-Membrane system: To generate the human plexin-B1 starting model (res. 1479-2135) for the simulations, structure of the Plexin-B1 intracellular region, was obtained from the Protein Data Bank (PDB ID: 3SU8)^17^ and the TM segment was N-terminally appended on the helical juxtamembrane segment as an ideal alpha-helix, however, leaving a small gap^26^. Such missing segments were reconstructed using Modeller^27^. After preparing the Plexin-B1 structure, the human Rnd1 and Rap1b GTPases were positioned onto the complex by structural superimposition, using available Plexin-B1-GTPase crystal structures as templates (PDB IDs: 2REX, 3SU8, and 4M8N)^17,28,29^. The bound GTP molecule and magnesium (Mg^2+^) ions for Rnd1 and Rap1b were subsequently aligned and transferred onto the modeled complexes based on the corresponding crystal structures. Prenylation of both GTPases at their C-termini was explicitly included. Rnd1 was modeled with a farnesyl modification at Cys229, whereas Rap1b carried a geranylgeranyl modification at Cys181. Force-field parameters for these lipid modifications were generated using Cgenff within the Charmm-Gui platform^30^ to be compatible with the all-atom CHARMM36m potential function, which is used throughout the project.

Following preparation of protein structures, membrane-embedded systems were constructed using the Charmm-gui webserver, membrane builder plugin^30,31^. The protein orientation of the TM region within the lipid bilayer and of the proteins interacting with the membrane peripherally was optimized and adjusted using the PPM(Positioning of proteins in membranes) module within the Charmm-gui webserver. All simulation systems consist of a mixed lipid bilayer containing 95% 1-palmitoyl-2-oleoyl-sn-glycero-3-phosphocholine (POPC) and 5% of PIP lipids – either Phosphatidylinositol 4,5-bisphosphate(PIP2) or Phosphatidylinositol 3,4,5-trisphosphate(PIP3) - on both upper and lower leaflets. We note here that in physiological cells an asymmetry of the bilayer is maintained by lipases, preferentially localizing the PIP2 (and other negatively charged lipids such as PS) on the cytoplasm facing leaflet of the membrane^32^. However, here we simulate a symmetric bilayer, which is typically seen in cancer cells. In any case, only the PIP2/PIP3 concentration would be slightly diminished in the outer leaflet, but since no elaborate extracellular segments of plexin are included in the simulation, no effect is expected.

A total of four systems were constructed: Plexin-B1_Rnd1_Rap1(B1_RnRp_PIP2) with PC/PIP2, Plexin-B1_Rnd1_Rap1b with PC/PIP3(B1_RnRp_PIP3), Plexin-B1_Rap1b with PC/PIP2(B1_Rp_PIP2) and Plexin-B1_Rap1b with PC/PIP3 (B1_Rp_PIP3). Each system was solvated with TIP3P water molecules^33,34^, neutralized, and further Na^+^/CL^-^ ions were added to give a concentration of 150mM at physiological pH conditions. A pH of 7.0 was used, nominally setting HIS residues to the neutral HSD isoform. Systems with Plexin-B1 and both GTPases were built in a cubic simulation box of 175 Å x175 Å x 200 Å dimensions and contained around ∼550,000 atoms, whereas systems with Plexin-B1 and Rap1b included ∼500,000 atoms.

Molecular Dynamics Simulation: Each system was energy minimized, which was then followed by equilibration, treated as a NPT ensemble (at 1bar and 300K). As per Charmm-gui, the standard six-step equilibration protocol was applied, in which, in each step, position restraints of the atoms were gradually released at each step. In order to remove boundary effects, periodic boundary conditions (PBC) were used. Nonbonded interactions were treated using a 12 Å cutoff for van der Waals forces, while long-range electrostatics were computed using the Particle Mesh Ewald (PME) method^35^. All covalent bonds involving hydrogen atoms were constrained using the LINCS algorithm^36^. The equilibrated systems were converted to Desmond format for Anton3 simulation^37^. Three independent 1μs production simulations for each system were performed (a total of 12 simulations each of 1μs) at 310 K and 1 bar on the Anton 3 supercomputer maintained by the Pittsburgh Supercomputing Center (PSC) and provided by D. E. Shaw Research.

Analysis of Trajectories: Protein-membrane interactions were calculated using in-house TCL scripts, in which the interaction frequency of each residue with individual lipid type (PC/PI) was computed using a 3.6Å atom to atom cutoff. Residues close to each other greater than 20% of the time and present across at least 2 of 3 replicas were filtered for visualizations. Membrane-protein contact areas were computed using the *measure sasa* function within VMD^38^, with customized TCL scripts adapted for this calculation. The orientation (angle) between the juxta-membrane helical region (1519 to 1549) with respect to the membrane and the distance between protein and bilayer were calculated with in-house scripts. For contact map analysis, we first performed cluster analysis using *gmx_cluster* within Gromacs by combining trajectories from all replicas for each system (3 μs per system) and using a cutoff of 0.6-0.7 Å. The representative structure from the major cluster was then used as input for the *Mapiya contact web server*^39^ to generate a residue-residue interaction dataset (with a cutoff of 5.0 Å). All plots were generated in R (version 4.2.0), and structural visualizations were produced using VMD^38^ (version 1.9.4). For network analysis, we followed the same protocol as used in our previous work, i.e., using the NAPS webserver^25,40^ (see detailed explanation in Bhattarai et al., 2025^21^). Local frustration analysis was performed using the Frustratometer package^41^ following the same approach as used by Bhattarai et al^25^, as well.

## Results

### 1.1 Protein-Membrane Localization

To assess how well the proteins are stabilized with the lipid bilayer, we first visualized all the complexes. As shown in Figure 1b, in the initial structure, the proteins are positioned well above the membrane, with lipidated C-termini of both GTPases within the bilayer and the transmembrane region embedded (Figure 1c). Figures 1d-1g present snapshots for each system at the end of the simulations, where each protein is colored differently and interacting lipids are shown within 3.5 Å. Across all complexes, GTPases (Rnd1/Rap1) remained stabilized with the membrane, and several regions in Plexin-B1 also formed consistent interactions with the bilayer.

To quantify the membrane localization of each complex, we calculated the center-of-mass (COM) distance along the Z-axis between the Cα atoms of the protein complex and the phosphate atoms of the lipid headgroups for all replicas in every system. For each complex, we then plotted the average COM distance across all replicas as a function of time (Figure 1h). At the start of the simulations, all systems showed an initial protein-membrane separation of ∼75 Å, reflecting their starting position above the bilayer surface. As the simulations progressed, this distance steadily decreased as the proteins approached and interacted with the membrane. At around 300 ns, most complexes had reached a stable membrane association, except for the B1_RnRp_PIP2/PIP3 system, which achieved a stable membrane-embedded state at roughly 500 ns. Furthermore, structural visualization (Figures 2a and 2b) reveals that Rnd1 and Rap1b engage slightly more extensively with the membrane in the PIP2 system (Figure 2), indicating the number and dynamics of lipid–residue interactions might differ between the two systems.

**Figure 2:**
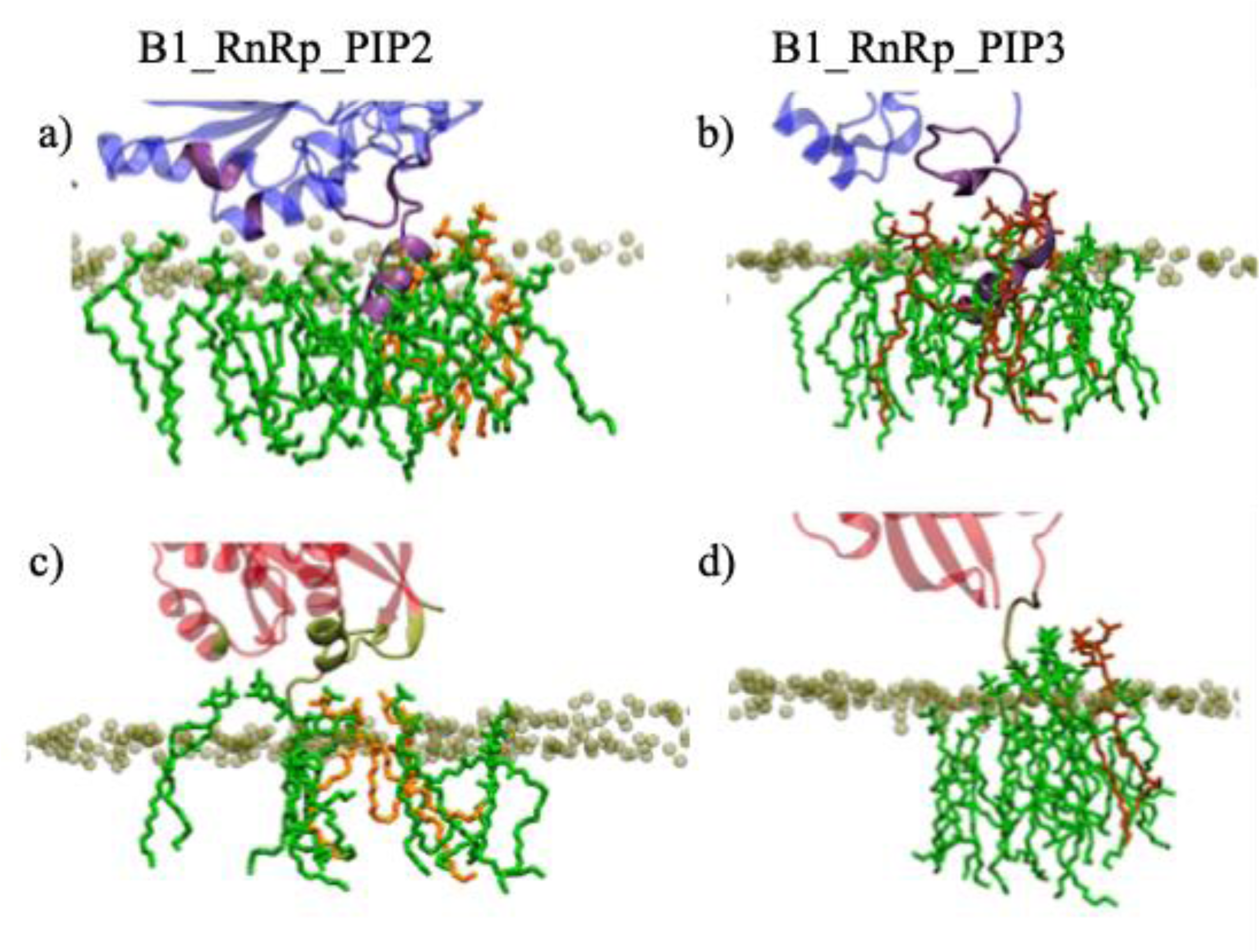
a-b) Visualization of Rnd1-membrane interacting region (colored in purple) for B1_RnRp_PIP2 and B1_RnRp_PIP3 systems, respectively. Some interacting PI/PC lipids are also shown. c-d) Visualization of Rap1-membrane interacting region (colored in olive) for B1_RnRp_PIP2 and B1_RnRp_PIP3 systems, respectively.

### GTPases-Lipid Interaction

We characterized residue-level lipid interactions with GTPases. To achieve this, we calculated the frequency of contacts between individual residues and lipid types (POPC and PIP2/PIP3) for Rnd1 and Rap1b across all simulation replicas (cutoff: 3.6 Å; total simulation time of 1.0 μs per replica). The interaction frequency indicates how often each residue interacts with a given lipid type (PC/PIP) during the entire simulation time. For each system, we filter for residues interacting with lipids in at least 2 replicas and average their occupancy. Interestingly, all Rap1b residues interacting with lipids in the PIP3 system were also observed in the PIP2 system, indicating broader lipid engagement in the presence of PIP2.

Figure 3a and 3b show bar plots of residue-lipid interaction frequencies for Rap1b in the B1_RnRp_PIP2 and B1_RnRp_PIP3 systems, respectively. In both systems, the lipid interactions were dominated by residues within the hypervariable region (HVR: 167-181) with higher contact frequencies. However, for the PIP2 system, more residues of the HVR region were involved in lipid interactions (V171, P172, G173, A175). In addition to the HVR, residues from other regions of Rap1b, such as R2, D47, Q49, and R136, N139, and N140, also formed lipid contacts, primarily with PC lipids; however, most of these interactions occurred with lower occupancy, consistent with transient membrane contacts. This trend was also observed in systems lacking Rnd1 (B1_Rp_PIP2 and B1_Rn_PIP3, Figure 3c and 3d). In the B1_Rp_PIP2 system, additional residues, including E45, D47, Q50, K128, R136, S150, K151, N153, N155, and E156, showed low-occupancy interactions with PC, which were absent in the corresponding PIP3 system. The set of residues commonly interacting with lipids in both B1_Rp_PIP2 and B1_Rp_PIP3 systems was largely similar to those observed in the presence of Rnd1 (Figure 3a, 3b). Together, these results indicate that Rap1b-lipid interactions are driven by HVR, and this was true across PIP2 and PIP3 systems, and PIP2 systems involve residues outside of the HVR and show a broader membrane engagement.

**Figure 3:**
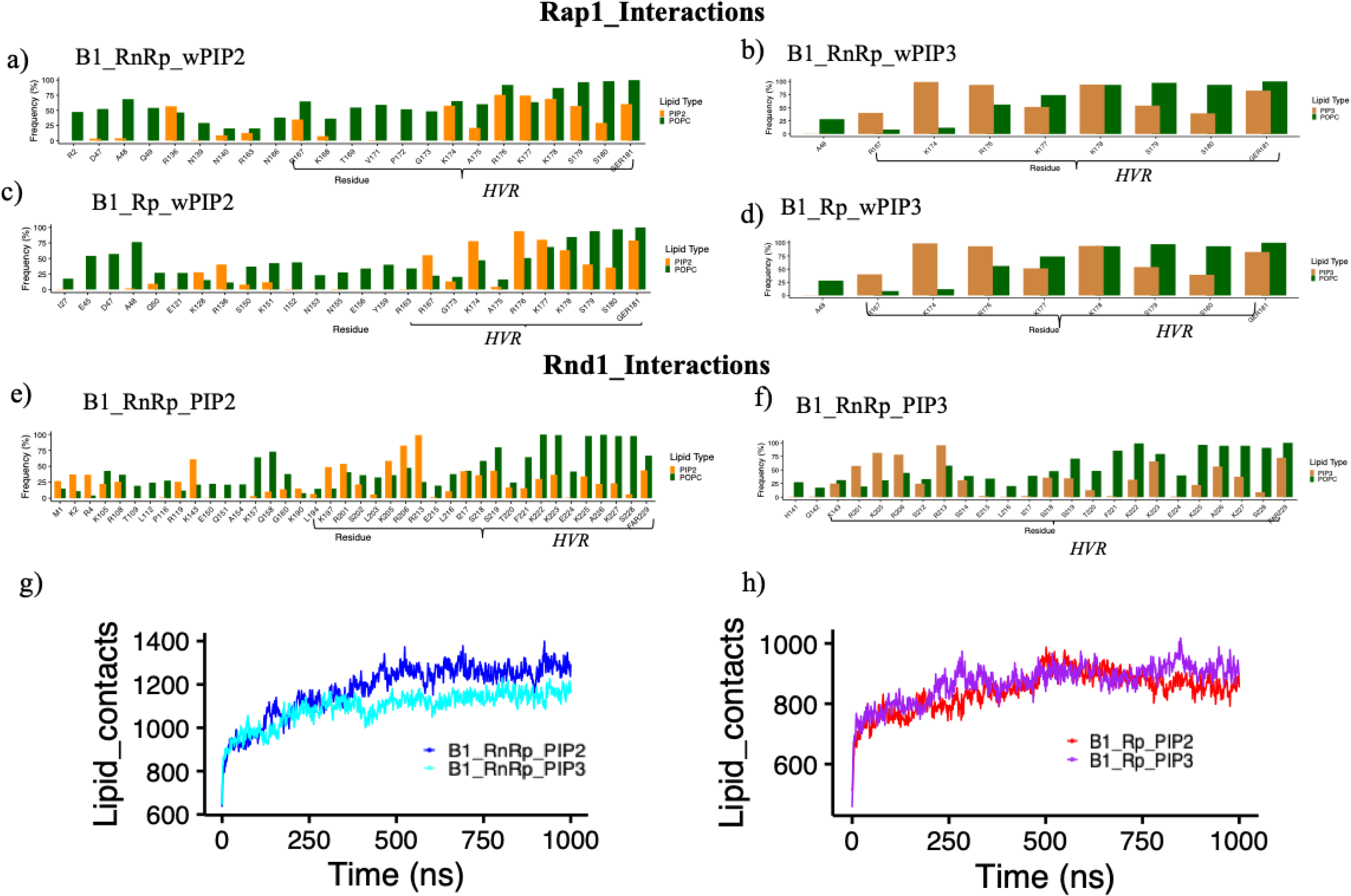
Bar plots showing the frequency of lipid interactions per residue, where phosphatidylcholine (PC) contacts are shown in green and phosphatidylinositol lipids PIP2 in orange and PIP3 in mustard color (a-d) with Rap1b (e-f) with Rnd1.Total number of lipid contacts between entire protein and lipid atoms (g) for B1_RnRp_PIP2/PIP3 systems h) for B1_Rp_PIP2/PIP3 systems

To access how residue-level interactions correlate to the overall total number of lipid-contacts, we quantified the total number of lipid-atoms contacting Rap1b using the same 3.6 Å cutoff. For this analysis, the total number of contacts between protein and POPC or PIP lipids was calculated separately for each replica, and all the data from the last 500 ns of the simulations were used to compare them (Supplemental Figure 1). With Rap1b, in the presence of Rnd1, Rap1b showed a higher number of contacts with POPC, consistent with the greater number of residues interacting with POPC (Supplemental Figure 1c and Figure 3a). Interestingly, despite more residues being involved, the total number of PI lipid contacts was slightly higher in the B1_RnRp_PIP3 system than in B1_RnRp_PIP2 (Supplemental Figure 1d). Opposite patterns were observed for Rap1b in systems without Rnd1 (Supplemental Figure 1e and 1f), with slightly more POPC contacts occurring in the B1_Rp_PIP3 system relative to B1_Rp_PIP2 and fewer PI contacts in the B1_Rp_PIP3 system. Overall, these results indicate that Rap1b-lipid interactions are both PIP-dependent and Rnd1-dependent, but Rnd1 exerts a larger influence, especially on non-PIP lipid contacts. PIP identity fine-tunes which residues engage the membrane, with PIP2 driving broader HVR interactions.

We further calculated residue-lipid interaction frequencies for Rnd1 GTPase and compared between systems between PIP2 and PIP3 (Figure 3e and 3f). Similar to Rap1b, a larger number of Rnd1 residues interacted in the PIP2 system. The majority of higher occupancy lipid interactions originated from the hypervariable region (residues: 180-229), with the PIP2 system, and additional HVR residues such as K190, L194, and K197 were also present. Beyond HVR, several residues from other regions of Rnd1, including K105, R106, K143, E150, A154, K157, and Q158, also interacted with lipids in PIP2 complexes with lower occupancy than HVR residues, suggesting transient and orientation-dependent lipid interactions. Analysis of the total number of contacts revealed (Supplemental Figure 1a and 1b) a slightly higher PIP contact in the PIP2 system, but not to an extent that would be considered significant. Together, these results indicate that Rnd1-lipid interactions are more favorable in the presence of PIP2.

Finally, comparison of total lipid contacts (PC + PIP) over the full simulation trajectory showed higher overall lipid engagement in Rnd1-containing PIP2 systems (Figure 3g, h). In contrast, in the absence of Rnd1, differences in total lipid contacts were minimal, suggesting that the lipid interaction differences observed for Rap1b (Figure 3b, d) are, at least in part, dependent on the presence of Rnd1. Overall, these findings suggest that PIP2-containing membranes promote stronger and more extensive GTPase–lipid interactions compared to PIP3-containing membranes. and lipid composition differentially modulates membrane engagement of Rap1b and Rnd1, with Rnd1 being more sensitive to PIP identity, while Rap1b membrane association appears to show some dependence on Rnd1 as well as PIP identity.

### 1.2 Plexin-B1 Interactions

After characterizing GTPase-membrane interactions, we next focused on Plexin-B1 membrane interaction and dynamics. For Plexin-B1, the primary membrane interacting regions include the transmembrane (TM), juxta-membrane (JM) region, and some sections of the activation switch loop (ASL) region. Figure 4a-4d shows bar plots of residue-level lipid interaction frequencies between Plexin-B1 and POPC or PIP2/PIP3 across all systems. For this computation, as well as the above, residues present in at least two replicas were considered.

**Figure 4:**
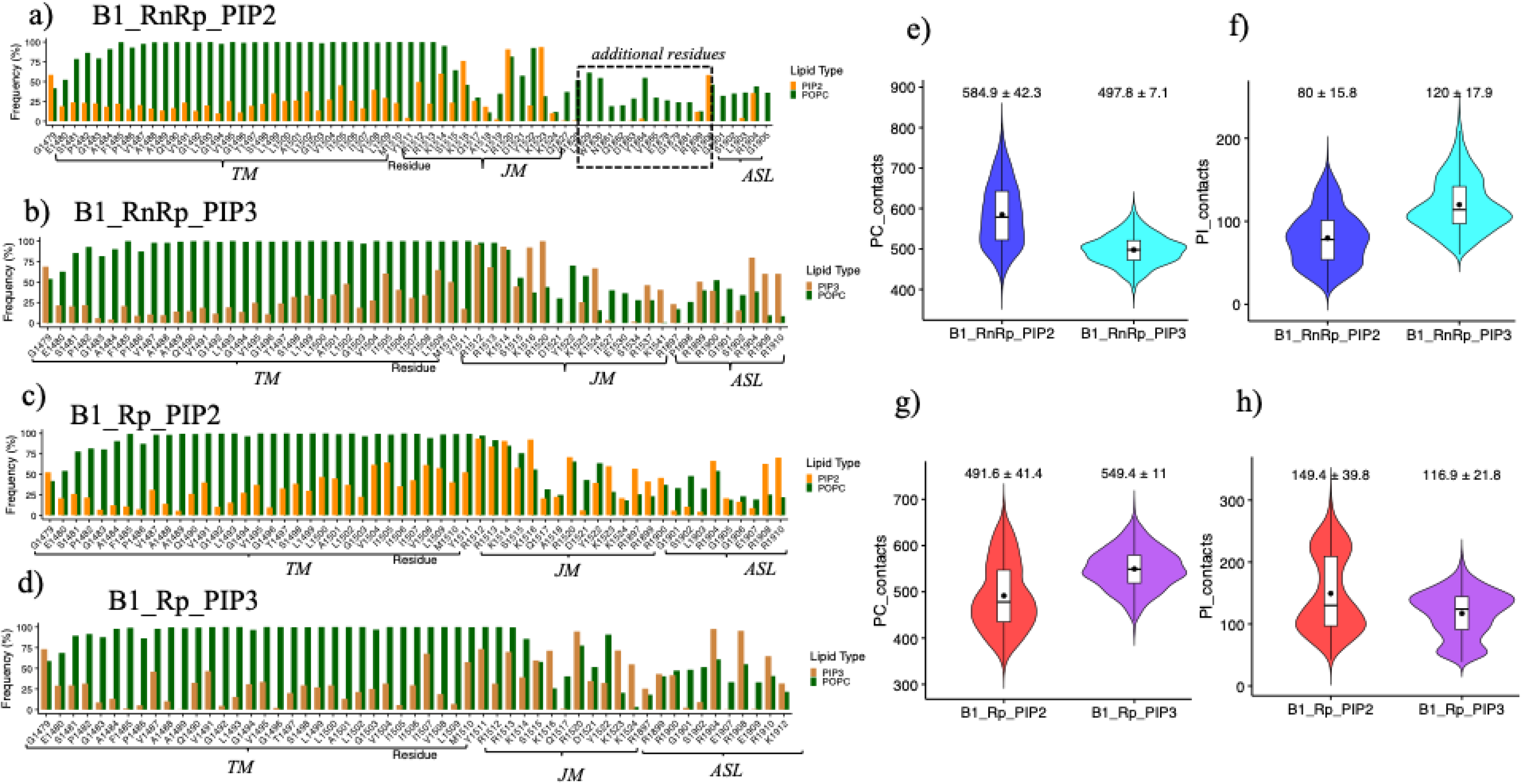
Lipid interactions with Plexin-B1.(a-d) Bar plots showing the frequency of lipid interactions per residue in Plexin-B1, with phosphatidylcholine (PC) contacts shown in green and phosphatidylinositol (PIP2/PIP3) contacts shown in orange and mustard color, respectively for all complexes. Violin plots depicting the total number of lipid contacts formed by Plexin-B1 in the *Plexin-B1_Rnd1_Rap1_PIP2/PIP3* systems over the final 500 ns of simulation (data from all three replicas shown): (e) PC contacts and (f) PI contacts. Violin plots illustrating the total number of lipid contacts formed by Plexin-B1 in the *Plexin-B1_Rap1_PIP2/PIP3* systems over the final 500 ns of simulation (data from all three replicas shown): (g) PC contacts and (h) PI contacts

Across all systems, the interacting residues predominantly localize to the transmembrane region: 1479-1511, juxta-membrane region: 1511-1545, and activation switch loop (ASL): 1890-1912. In the B1_RnRp_PIP2 system, additional cytoplasmic residues within the RBD and its linker region: G1828, L1829, W1830, N1861, Q1862, D1863, E1878, and G1879 also engage the membrane. These interactions were absent when either Rnd1 was missing or when PIP3 instead of PIP2 was present, indicating a cooperative, context-dependent membrane engagement that requires both Rnd1 and PIP2. As shown in Figure 3, Rnd1 and Rap1b also display enhanced lipid interactions in the B1_RnRP_PIP2 complex, suggesting that the B1_RnRp_PIP2 complex stabilizes a PlexinGAP/ASL orientation which facilitates additional RBD contacts with the membrane, while Rap1b engagement is also concurrently enhanced. Collectively, these results highlight a synergistic effect of protein-protein and protein-lipid interactions, driven by PIP2 and the presence of both binding GTPases. Our previous study reported that Plexin-B receptor activation switch loop that controls its GAP activity toward Rap1b in a cooperative and kinetically allosteric and substrate-dependent manner^9^.

Some additional interactions with the ASL region, residues R1908 and R1910, which are present in other systems, are absent in the B1_RnRp_PIP2, indicating structural differences in Plexin-B1 membrane-interacting regions between systems. In addition, for systems without Rnd1, additional residues E1907 and R1908 were also interacting with PIP lipids. Structural visualization further reveals distinct conformational and dynamical differences between the B1_RnRp_PIP2 and B1_RnRp_PIP3 systems, where an additional loop region (labeled in Fig. 4a) in the PIP2 system engages with the bilayer, while the same region adopts a different conformation in the PIP3 system that precludes membrane interaction.

To further quantify residue-level interaction frequencies, we used the last 500 ns of the simulations across all replicas to compute the total number of contacts between Plexin-B1 and each lipid type. In the presence of Rnd1 in complex, although the B1_RnRP_PIP2 contains a greater number of interacting residues (Figure 4a), the total number of plexin-PI lipid contacts was a little higher in the B1_RnRp_PIP3 system than in the B1_RnRp_PIP2 (mean: 120 ± 18 vs. 80 ± 16; Figure 4f) and slightly more POPC contacts were formed for the B1_RnRp_PIP2 system (Figure 4e).

In the absence of Rnd1, an opposite trend was observed, with the plexin in B1_Rp_PIP3 system exhibiting slightly higher PC contacts (491 ± 41 vs 549 ± 11, Figure 4g) and the B1_Rp_PIP2 system showing increased PI contacts (149 ± 39 vs 117 ± 22, Figure 4h). This behavior was also observed for Rap1b in systems lacking Rnd1 (Figure 3i and 3j). It should also be noted that compared to the total number of lipid- protein atom interactions plotted in Fig. 3g & h, the total number of PIP lipid contacts was less, where the average from all three runs was ∼154 for B1_RnRp_PIP2 and ∼201 for B1_RnRP_PIP3 systems. The number of PIP2/3 contacts is relatively small with dually complexed plexin, whereas with plexin with Rap1b alone, the total contacts for PIP lipids were ∼145 in the PIP3 system and increased contacts to ∼165 in the PIP2 system, suggesting that at least some of the non-Rnd1 interacting surface is now available for interactions. Remarkably, the number of plexin-PIP3 contacts remains the same, whether Rnd1 is present or not, emphasizing again that there is lipid isoform specificity to interactions. Together, these results indicate that Plexin-B1 membrane engagement is modulated by both lipid composition and GTPase context. While lipid identity influences which residues interact and the type of lipid contacts, Rnd1 and Rap1b stabilize Plexin-B1 conformations and promote synergistic membrane interactions.

### 1.3 Juxta-Membrane Segment – Membrane interactions

During the 1.0 μs simulations, moderate structural changes were observed in multiple protein regions, specifically in the Plexin-B1 juxta membrane (JM) region relative to the GAP/RBD region. These changes appear to be influenced by both lipid composition and associated GTPases. To quantify these effects, we calculated the time evolution of the angle between bilayer normal (defined by phosphate atoms in the upper leaflet) and the juxta-membrane helical region (residues: 1519 to 1549) for all replicas and systems (Figure 5a). We then plotted the average with simulation time (Figure 5b and 5c), showing that most of the reorientation change is complete by 500 ns,. To analyze the stable conformation following structural rearrangement, the final 500 ns data of each simulation were used to compute box plots confirming this behavior(Inset: Figure 5b and 5c).

**Figure 5:**
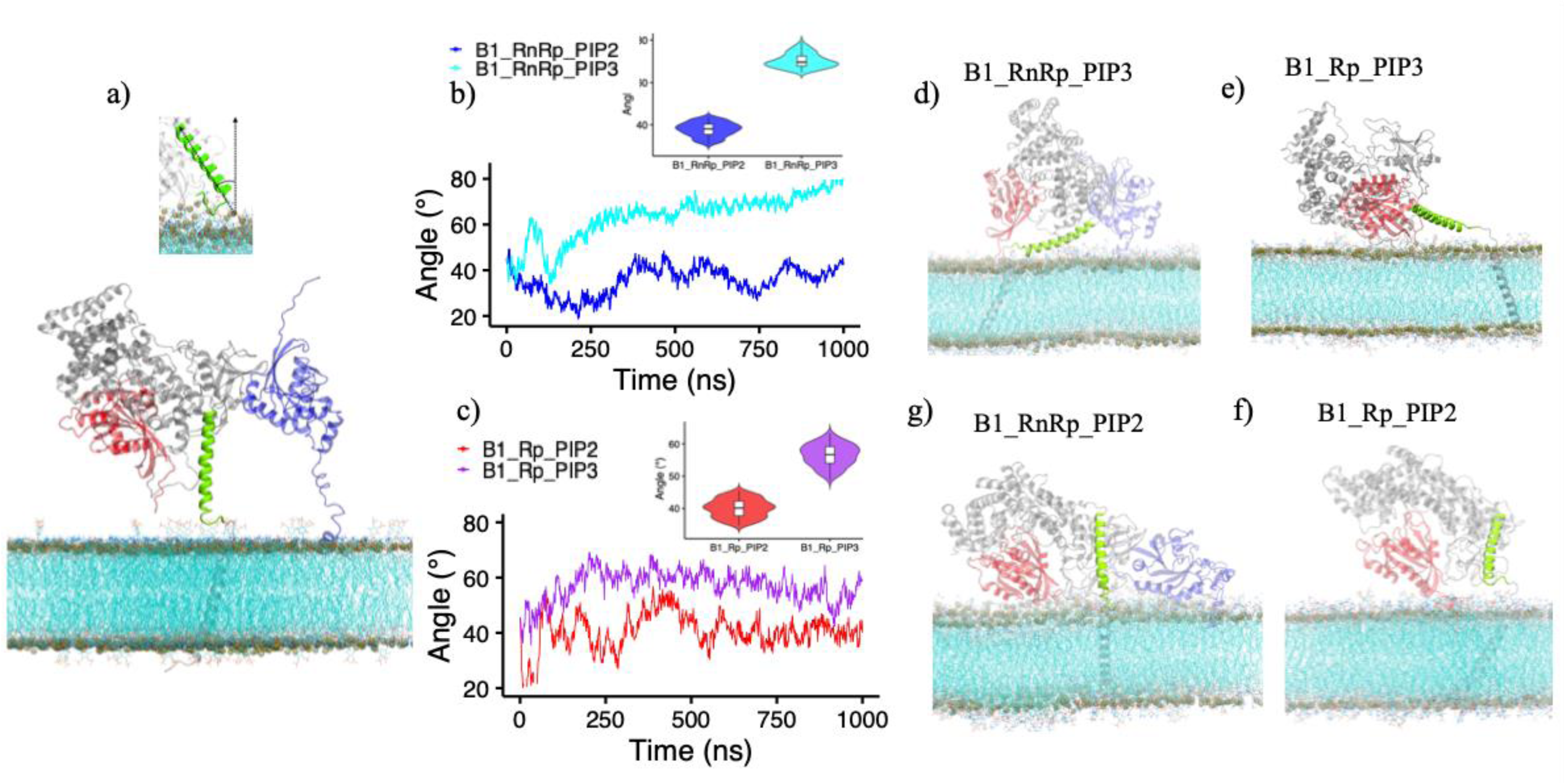
Juxta-Membrane Dynamics a) Initial system configuration highlighting juxta membrane region in green(resid: 1519 to 1549), Plexin-B1 in gray, Rap1b in red, and Rnd1 in blue. Time evolution of the angle between the normal of phosphate atoms at the upper leaflet of the membrane and JM b) for B1_RnRp_PIP2/PIp3 systems, c) for B1_Rp_PIP2/PIP3 systems. Insets in both panels show box plots of the angle distributions over the last 500 ns.(d-g) Snapshots highlighting JM orientation (colored green) for all complexes around the end of 1us simulation time.

The B1_RnRp_PIP3 system (containing both Rnd1 and PIP3) shows the largest angular deviation of the JM region (mean: 70.3° ± 3.4°), suggesting substantial structural changes. In contrast, the B1_RnRp_PIP2 system shows a significantly lower mean angle (37.7° ± 3.3°) (Fig. 5b). Interestingly, in the absence of Rnd1, a similar difference is observed, though with reduced magnitude (mean JM angle is 56° ± 3.4° for B1_Rp_PIP3 and 40.1° ± 2.8° for B1_Rp_PIP2, Fig. 5c).

Visualization of the JM regions further supports these findings, revealing that in PIP3-containing systems the JM helix deviates most strongly and adopts nearly parallel orientation to the bilayer, whereas in PIP2-containing systems, the JM region remains predominantly vertical regardless of the presence of Rnd1. (Figure 5e and 5g). While the JM region undergoes the greatest reorientation in the B1_RnRp_PIP3 system, overall lipid interactions are higher in the B1_RnRp_PIP2 system, indicating that JM reorientation may play a limited role in membrane interactions and could be associated with other functional effects. These results indicate that Plexin-B1 JM orientation is strongly modulated by lipid composition, with PIP3 promoting pronounced JM reorientation toward a horizontal membrane-aligned state, an effect further enhanced by Rnd1. The JM region is an interaction partner with the ASL holding the loop away from the GAP region which allows for faster substrate entry and product Rap1b exit^11^. The finding that with PIP3, this interaction is diminished suggests that PIP3 could be activating plexin’s GAP function.

### 1.4 Interaction between Plexin-B1 and GTPases

The interaction between Plexin-B1 and Rho GTPases is thought to play a role in inactivating or activation Plexin-B1’s GAP activity for helping to hydrolyse he GPT nucleotide in the Ras family Rap1b GTPase. In our previous study, we reported that the type of GTPase bound to Plexin-B1 in solution influences allosteric communication across the protein, modulating its interaction network^21^. Here, we placed the simulation system at a model membrane and explored how lipid composition affects Plexin-Rap1b interaction and the presence or absence of Rnd1.

To assess this, we generated a contact map between Rnd1/Rap1b and Plexin-B1. For the contact map, cluster analysis was performed combining all three replicas (total simulation time : 3 μs), and the dominant cluster, which in each system represented at least 50-60% of the conformations was used as an input to the Mapiya webserver to extract the contact maps as Figure 6. Figure 6a and 6b show the contact maps between Rnd1 and Plexin-B1 for the B1_RnRp_PIP2 and B1_RnRp_PIP3 systems, respectively. In both systems, most interactions of plexin’s GAP occur with residues in the RBD region. Interestingly, the PIP3-containing system exhibited more Rnd1-Plexin-B1 (total: 48 and stronger) contacts compared to the PIP2 system (total: 40), suggesting that lipid composition influences the extent of these interactions.

**Figure 6:**
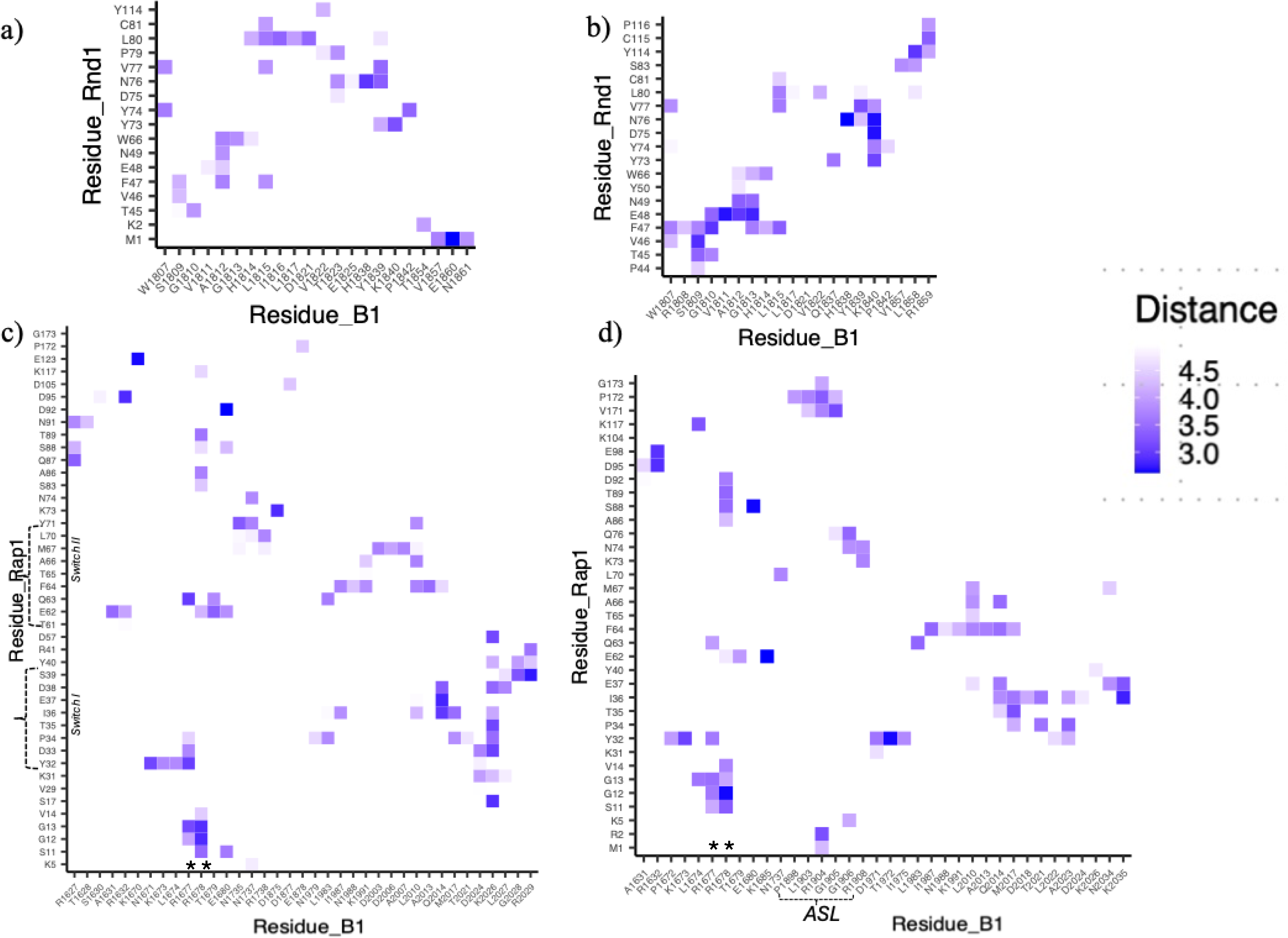
Contact map showing interactions between GTPases and Plexin-B1 a) between Rnd1 and Plexin-B1 in Plexin-B1_Rnd1_Rap1 with PIP2 system b) between Rnd1 and Plexin-B1 in Plexin-B1_Rnd1_Rap1 with PIP3 system c) between Rap1 and Plexin-B1 in Plexin-B1_Rnd1_Rap1 with PIP2 system d) between Rap1 and Plexin-B1 in Plexin-B1_Rnd1_Rap1 with PIP3 system.(**-indicates arginine finger region)

However, Rap1b showed more connections with Plexin-B1 in the B1_RnRp_PIP2 system: a total of 97, compared to 79 connections in the B1_RnRp_PIP3 system. These additional interactions involve regions outside the Rap1b switch regions. Interestingly, we observe an engagement of ASL residues in B1_RnRp_PIP3 complexes with lipids (Figure 4d). These results indicate that lipid composition may modulate interactions between Plexin and GTPase. Notably, the B1_RnRp_PIP3 system also exhibited the largest JM angle deviation and reduced Rnd1-lipid interactions, suggesting that structural rearrangements may facilitate Rnd1-Plexin-B1 contacts. Similarly, Rap1b-lipid interactions were slightly stronger in the PIP2-containing systems (Figure 3), which might have led to weaker interactions between Plexin-B1 and Rap1b. In our previous study of plexin GTPase complexes in solution^25^, a stronger Rap1b-plexin B1 interaction network was observed in the absence of Rnd1. A similar trend is observed in the membrane-bound systems, where Rap1b-B1 connectivity appears to depend more strongly on the presence of Rnd1 than on PIP identity.

We also examined protein-protein contacts in the systems lacking Rnd1 (Supplemental Figure 2). Here, Rap1b formed more connections with Plexin-B1 in the PIP3 system (total: 84) compared to the PIP2-containing system (total: 74), reflecting differences between plexin-GTPase and protein-lipid interactions in the PIP2 system. These results suggest that Rap1b dynamics and interactions with Plexin-B1 are more dependent on the presence of Rnd1 than on the lipid composition itself: in the presence of Rnd1, Rap1b forms more extensive contacts with Plexin-B1 in PIP2-containing systems.

We next assessed the network communication and dynamics between plexin-B1 and GTPases by integrating network connections and frustration analysis, using the same cluster center representative structures as above. The methodologies for network construction, significance, and frustration calculations have been described in detail in our previous work^21^ and in the methods section. Betweenness centrality quantifies the number of shortest communication paths passing through a residue, where higher values indicate residues that act as communication hubs or bridges. In the frustration analysis highly frustrated regions often correspond to functional or allosteric sites, whereas minimally frustrated regions typically stabilize the folded core.

We first visualized network connections, focusing on the plexin-B1 and GTPases interface, where we did not observe significant changes between plexin-B1 and Rnd1/Rap1b connections and PIP2/3 lipid type (Supplemental Figures 3 and 4). However, we did observe some changes when plotting network centrality. Figure 7a and 7b show residue-wise betweenness centrality for Rnd1 and Rap1b, respectively, for the B1_RnRp_PIP2 and B1_RnRp_PIP3 systems. For both GTPases, PIP3-containing systems showed regions with slightly higher betweenness for both GTPases, especially in switch regions, suggesting more efficient communication pathways in the presence of PIP3. For systems without Rnd1, we did not observe significant changes at the residue-level betweenness (Supplemental Figure 4a).

**Figure 7.**
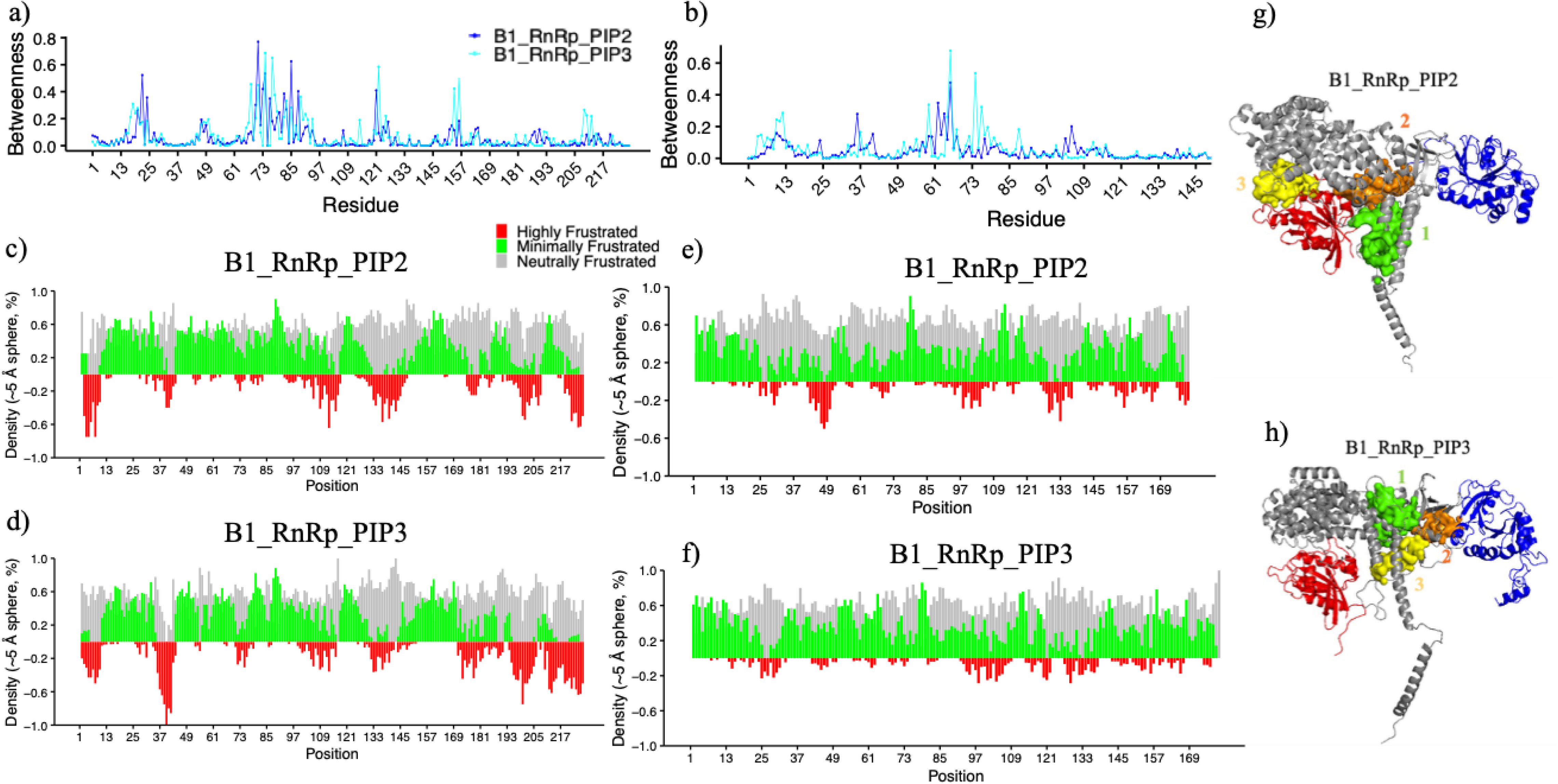
Network and frustration analysis. (a–b) Residue-wise betweenness centrality profiles for (a) Rnd1 and (b) Rap1b in the B1_RnRp_PIP2 (blue) and B1_RnRp_PIP3 (cyan) systems.(c–f) Local frustration analysis of (c) Rnd1 in B1_RnRp_PIP2, (d) Rnd1 in B1_RnRp_PIP3, (e) Rap1b in B1_RnRp_PIP2, and (f) Rap1b in B1_RnRp_PIP3.(g–h) Predicted allosteric site pockets identified on the major cluster structures in (g) B1_RnRp_PIP2 and (h) B1_RnRp_PIP3, with the top three pockets highlighted in distinct colors.

These results also correlate with local frustration analysis (Figures 7c-7f). Rnd1 displays regions enriched in highly frustrated interactions in both lipid environments, with a slightly higher prevalence in the PIP3 system (Figure 7d compared to PIP2 in Figure 7d), indicating allosteric or functional potential at the Plexin-B1-Rnd1 interface. For Rap1b as well, we observe regions that were highly frustrated for both systems; however, the extent of high frustration was slightly more in PIP2 systems (Figure 7e and 7f). For systems without Rnd1 as well, we observe several highly frustrated regions as well, slightly pronounced with B1_Rp_PIP2 systems (Supplemental Figure 5). However, overall frustration in Rap1b is higher in the absence of Rnd1, than in its presence. For Plexin-B1, across all systems, the transmembrane (TM) and juxtamembrane (JM) regions consistently exhibit the highest levels of local frustration, suggesting that these regions retain substantial conformational flexibility regardless of lipid composition or Rnd1 binding. In contrast, the activation switch loop (ASL) was largely frustrated in the B1_RnRp_PIP2 system(Supplemental Figure 6).

To further explore potential functional hotspots, we employed the PASSER webserver^42^ to visualize the top three allosteric site pockets using the major cluster. Figures 7g and 7h depict predicted allosteric pockets for B1_RnRP_PIP2 and B1_RnRP_PIP3 systems, respectively, with the top three pockets highlighted in distinct colors and labeled as 1, 2, and 3. Interestingly, in the B1_RnRp_PIP2 systems but also in the two systems without Rnd1, the top three pockets were located at or near the plexin-B1-Rap1b interface, which is also the region where arginine fingers R1677 and R1678 are located, with around 50% probability for all. In contrast, for the B1_RnRp_PIP3 system, the top three sites were at the Plexin-B1-Rnd1 interface or near the RBD-RND1 region, with around 50% probability. This, together with the data above, suggests that in the presence of Rnd1 and PIP3, the packing of Rap1b into the GAP domain may be more optimal (and thus have higher activity) than in the other cases.

## Discussion and Conclusion

Plexins are single-pass transmembrane receptors for Semaphorins that interact with Rho-family GTPases (Rnd1, Rac1, RhoD) and Ras-family members such as Rap1b. These two types of GTPases bind distinct regions of Plexin-B1, mainly the RBD and GAP domains, respectively, modulating its conformation and signaling activity. Because GTPase orientation and membrane localization can influence plexin–GTPase interactions and their allosteric communication, the lipid environment is likely to play an important regulatory role. While previous structural and biochemical studies have established that Rnd1 and Rap1b regulate Plexin activation through distinct interactions with the intracellular region, how these interactions are modulated by the surrounding membrane has remained unclear. Here we show by computational modeling and all-atom dynamics simulation that membrane composition plays an important role in determining the orientation and membrane engagement of lipid-modified small GTPases and their associated signaling complexes. In this study, we investigated how membrane composition, specifically PIP2 versus PIP3, affects Plexin-B1 conformational dynamics, membrane association, and allosteric coupling with GTPases. We also examined whether Rnd1 binding at the RBD alters these effects. To this end, we performed 1μs simulations in three replicas of four systems.

Examining the results, all systems were well localized to the membrane as depicted by largely plateaued values for the center of mass distance between protein and membrane(Figure 1h). At the residue level, Rnd1 and Rap1b GTPases maintained stable interactions with the lipid bilayer across all systems, with stronger membrane association in PIP2-containing membranes. In particular, PIP2-containing membranes promoted broader membrane engagement and involvement of a larger number of hypervariable region (HVR) residues for both GTPases. This behavior is consistent with previous studies showing that the membrane orientation and dynamics of lipid-modified small GTPases are governed by electrostatic interactions and anionic lipids. For example, work by Prakash P. and A. Gorfe demonstrated that the HVR and adjacent basic residues determine how GTPases sample membrane-bound orientations and interact with lipid headgroups. We have shown a similar orientational preference of KRas on lipid composition experimentally and also computationally for KRas4b^21,22^. Here, our results extend these observations to the Plexin-B1 signaling system, showing that phosphoinositide identity can subtly reshape how Rap1b and Rnd1 engage the membrane and interact with their receptor partner.

For Plexin-B1, the majority of lipid interactions were formed between residues at transmembrane and juxtamembrane (JM) regions and for activation switch loop (ASL) regions, with lipid composition modulating both the number of contacts and the specific residues involved (Figure 5). We also found that in PIP3-containing systems, the JM region adopted a more parallel orientation relative to the bilayer, particularly in the presence of Rnd1, whereas in PIP2 systems, the JM helix remained largely vertical (Figure 6). These results suggested phosphoinositide identity can influence the structural dynamics of membrane-proximal regions, potentially impacting signal transmission across the membrane. Because the JM region interacts with the activation switch loop that regulates accessibility of the GAP domain^11^, such lipid-dependent structural changes may influence Plexin-B1 activation and its regulation of Rap1b. These observations are consistent with structural and mechanistic studies showing that conformational changes within the cytoplasmic region regulate the level of plexin activity^11,13^.

The GTPase-receptor interface further supports a model of lipid-dependent allosteric redistribution rather than uniform strengthening or weakening of binding. Upon investigating the Plexin-B1 GTPase interface, contact map analysis demonstrated that Plexin-B1-GTPase interactions are sensitive to both lipid composition and Rnd1 presence. Rnd1-Plexin-B1 contacts were more extensive in PIP3 systems, correlating with JM reorientation and reduced Rnd1-membrane interactions (Figure 7). Similarly, for the Rap1b-Plexin-B1 interface, contacts were stronger in PIP2-containing membranes when Rnd1 was present, also indicating cooperative modulation of GTPase binding. These findings align with previous computational studies showing that GTPase binding can propagate allosteric signals across the Plexin cytoplasmic region and modulate GAP activity (Bhattarai et al., 2025)^25^. More broadly, they support emerging views that Ras- and Rho-family GTPases regulate their effectors through dynamic allosteric coupling and membrane-dependent conformational changes (Bhattarai & Buck, 2026)^24^.

To further understand the dynamics and allosteric sites between Plexin-B1 and GTPases, we also performed a network and local frustration analysis focusing on betweenness centrality. We found that systems/regions with high betweenness centrality often coincided with highly frustrated regions, representing potential allosteric hotspots. PIP2-containing systems favored functional frustration at the Plexin-B1-Rap1b interface, whereas PIP3 shifted these regions toward the Plexin-B1-Rnd1 interface. Again, predicted allosteric sites correspond with these frustrated regions, demonstrating that phosphoinositide identity can affect functional sites within the complex. Remarkably, Plexin-B1 in presence of PIP3 and Rnd1 was the least frustrated protein in the catalytic Arg finger region and also showed the tightest packing. This would suggest that activity may be highest of plexin-B1 at a PIP3 containing membrane in presence of Rnd1. No experimental studies have been reported to date of PIP2 or PIP3 on plexin activity. However, in other systems PIP2 is thought to be inhibitory, both in EGFR2 and EphA2^43^, **a**nd its down-regulation in cancers, increasing the PIP3/PIP2 ratio, indicates that PIP3 could be receptor activating. Experimental work will need to be carried out to substantiate this prediction, also, the experimental observation and computational suggestion that Rnd1 is activating the receptor, whereas RhoD and Rac1 are not^13,25,44,45^. Importantly, our findings also suggest specific experimental tests. To directly probe the JM–ASL coupling mechanism, mutations that disrupt electrostatic and structural communication between these regions would be particularly informative. For example, charge-neutralizing mutations of conserved lysine/arginine residues in the JM region (e.g., K→A or R→A substitutions in the membrane-proximal JM segment) could test the role of membrane anchoring in controlling ASL positioning.

Overall, our simulations reveal that lipid composition and GTPase context together shape the structural dynamics, membrane engagement, and communication pathways within Plexin-B1 signaling complexes. PIP2-containing membranes promote broader GTPase–lipid interactions and stronger Rap1b engagement with Plexin-B1, whereas PIP3 favors increased Rnd1–Plexin interactions and pronounced reorientation of the juxtamembrane region. These results highlight the membrane environment as an active regulator of Plexin signaling and suggest that phosphoinositide composition may influence how Plexin-B1 integrates signals from Rho- and Ras-family GTPases to control downstream cellular responses. Experimental tests of the predictions made in this computational study could involve studies with purified proteins, reconstituted at liposomes or in cells. Ideally, such research will be guided by the observations made here computationally; for example, one might expect the isoform specificity to be diminished if residues in the JM and ASL regions were mutated from Arg and Lys to other amino acids.

## Author Contributions

NB and MB designed the project. NB generated the protein coarse-grained models, performed the MD simulations, and analysed the data. ARS provided additional advice throughout. NB and MB co-wrote the paper. All authors read and approved the final version of the manuscript and supporting information. Initial stages of calculations on Plexin GTPase complexes in solution and at neutral membranes were also performed by previous postdoctoral fellows in Buck lab., Drs. ZhenLu Li and Liquin Zhang.

## Declaration of generative AI and AI-assisted technologies in the writing process

During the preparation of this manuscript, the authors used ChatGPT/GPT-5 to edit/improve the readability and language of the text. Following the use of this tool, the authors carefully reviewed and edited the manuscript as needed and accept full responsibility for the content of the published work.

## Acknowledgements

This work utilized the high-performance computing resources at CWRU and the Buck lab. It was supported by NIH grants R21AG084065, R01EY029169 when the project began and is currently supported by R01AG089561. NB was funded by a postdoctoral fellowship, NCI T32 (T32CA094186). We thank Pittsburg Supercomputing Center for providing computing resources (Anton3).

## Supporting Information Available

Supporting tables and figures are included in the supporting information

