## Supplemental Information for "A model for PIP2/3 and Rnd1 effects on Plexin-B1 GAP activity on Rap1b GTPase derived from molecular dynamics simulations"

**Supplemental Figure 1**

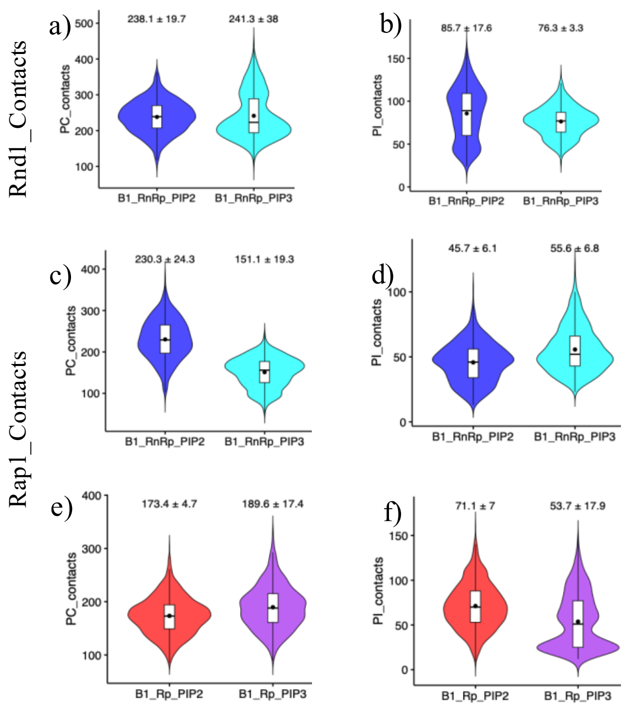


Violin plots depicting the total number of lipid contacts formed by Rnd1 in the *Plexin-B1_Rnd1_Rap1_PIP2/PIP3* systems: a) PC contacts b) PI contacts. Violin plots depicting the total number of lipid contacts formed by Rap1b in the *Plexin-B1_Rnd1_Rap1_PIP2/PIP3* systems c) PI-contacts d) PC contacts. Violin plots depicting the total number of lipid contacts formed by Rap1b in the *Plexin-B1_Rap1_PIP2/PIP3* systems e) PC contacts f) PI contacts. (Data from the last 500 ns across all replicas is shown. Mean with SEM is displayed for all plots.

**Supplemental Figure 2**

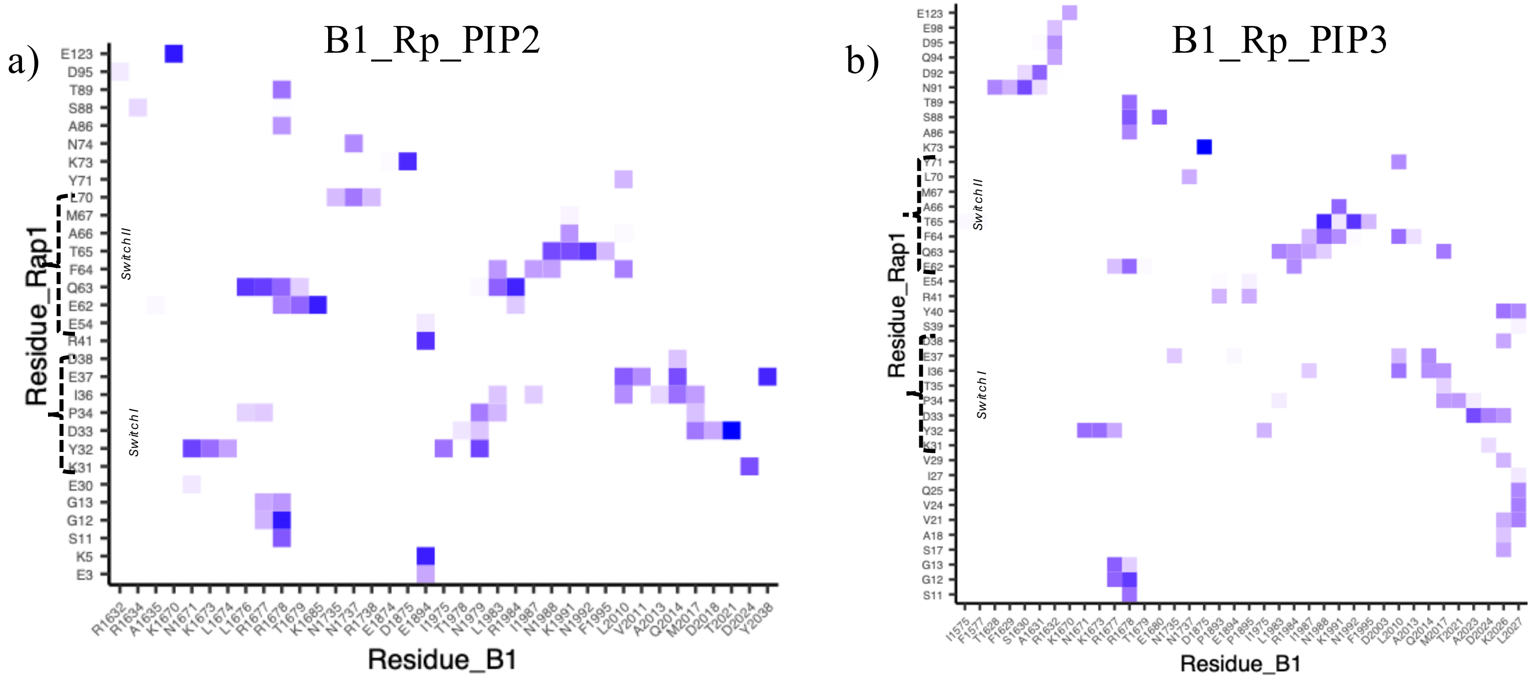


Contact map between Rap1b and Plexin-B1 a) For B1_Rp_PIP2 b) B1_Rp_PIP3 system

**Supplemental Figure 3**

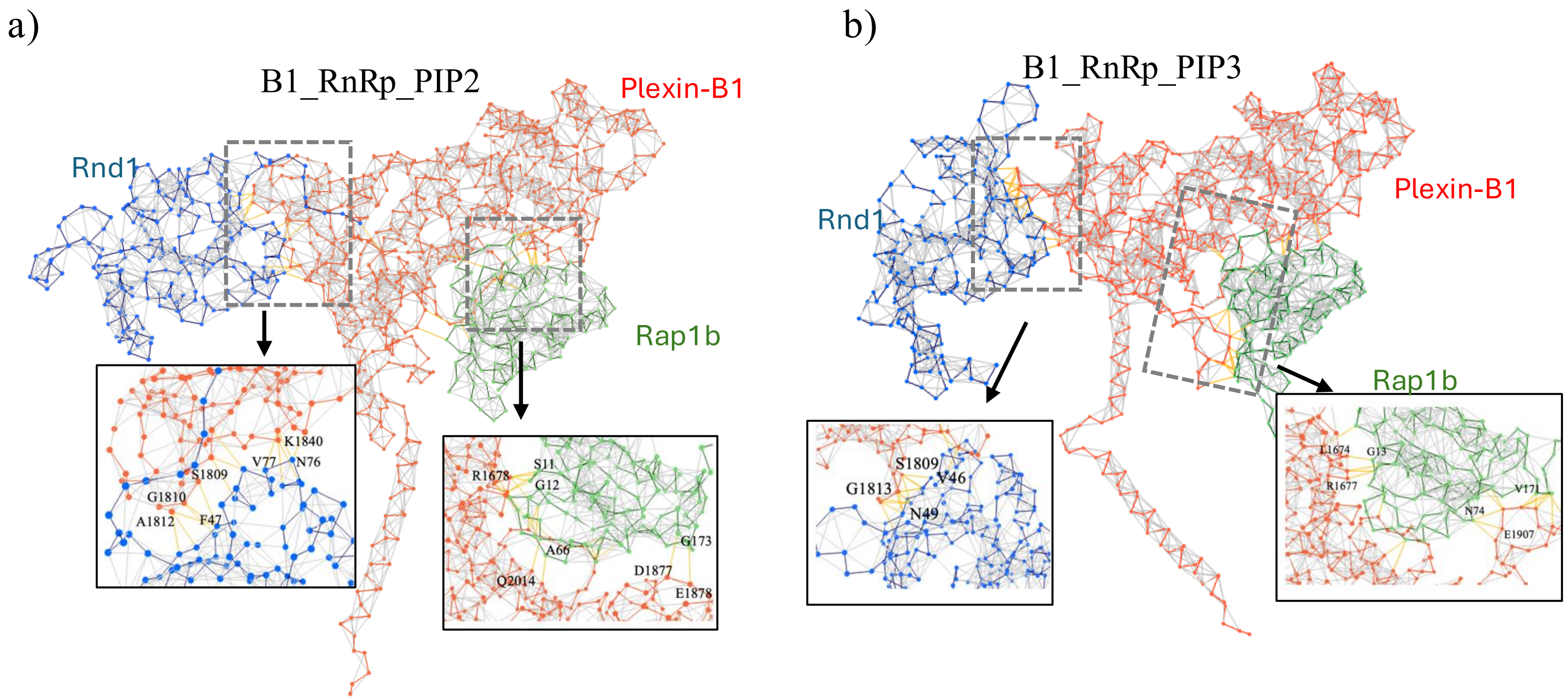


Network View from the major cluster a) For B1_RnRp_PIP2 b) B1_RnRp_PIP3 system, where Plexin-B1 is colored red, Rnd1 in blue, and Rap1b in green. The Zoom-in highlights connections between Plexin-B1 and GTPases, shown with yellow lines, and some residues involved in these connections are also indicated.

**Supplemental Figure 4**

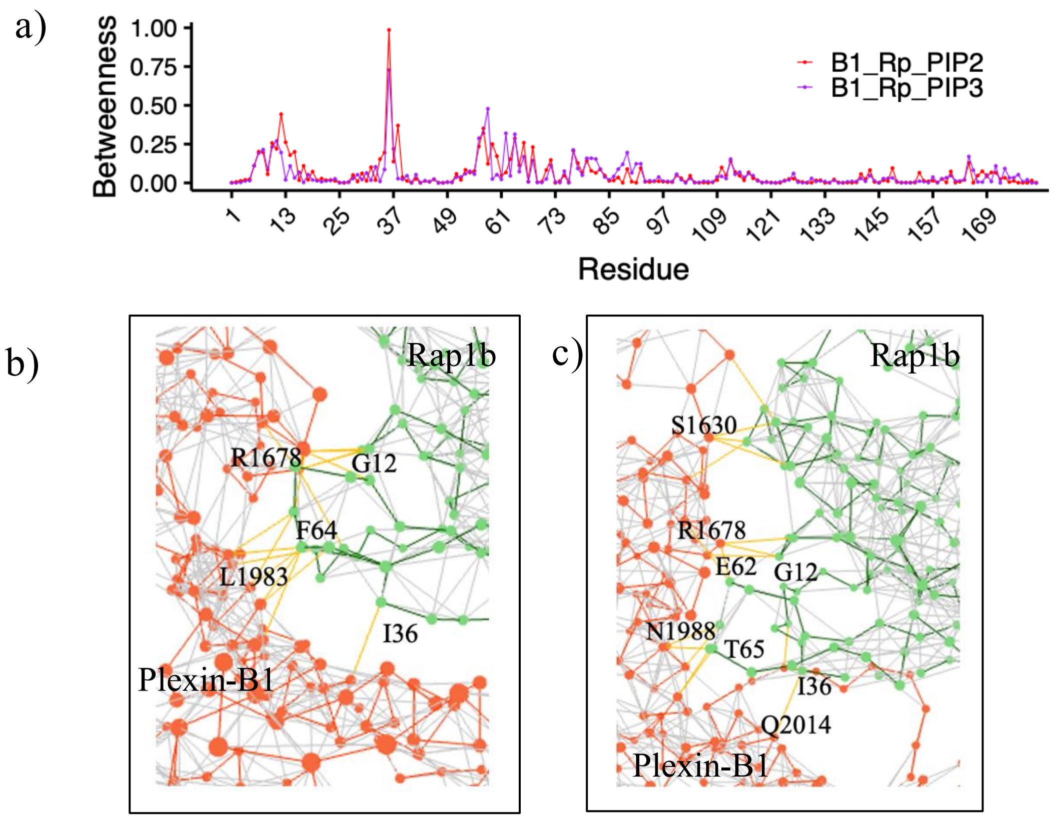


a) Residue-wise betweenness centrality for Rap1b in B1_Rp_PIP2 (colored red) and B1_Rp_PIP3 (colored purple) systems. b-c) Network connections between Plexin-B1 and Rap1b b) B1_Rp_PIP2 and c) B1_Rp_PIP3, shown with yellow lines, and some residues involved in these connections are also indicated.

**Supplemental Figure 5**

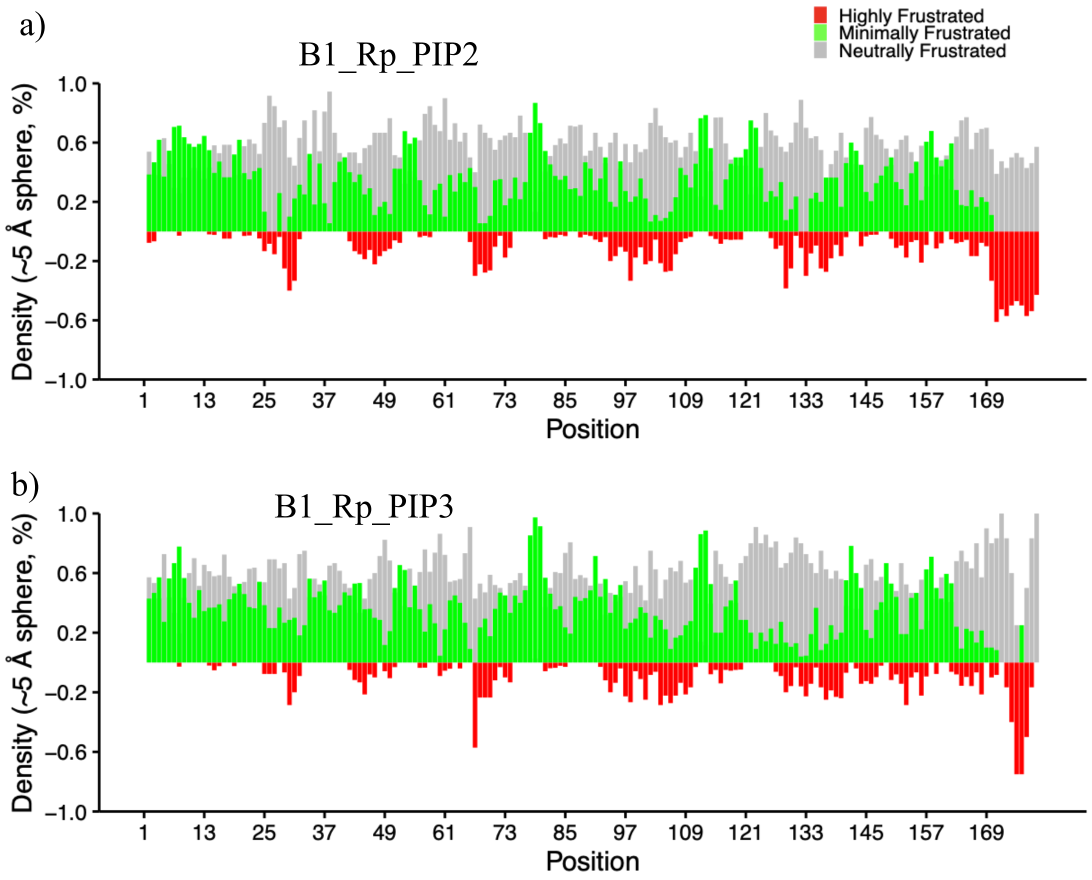


Local frustration analysis of Rap1b in (a) B1_Rp_PIP2, (b) B1_Rp_PIP3

**Supplemental Figure 6**

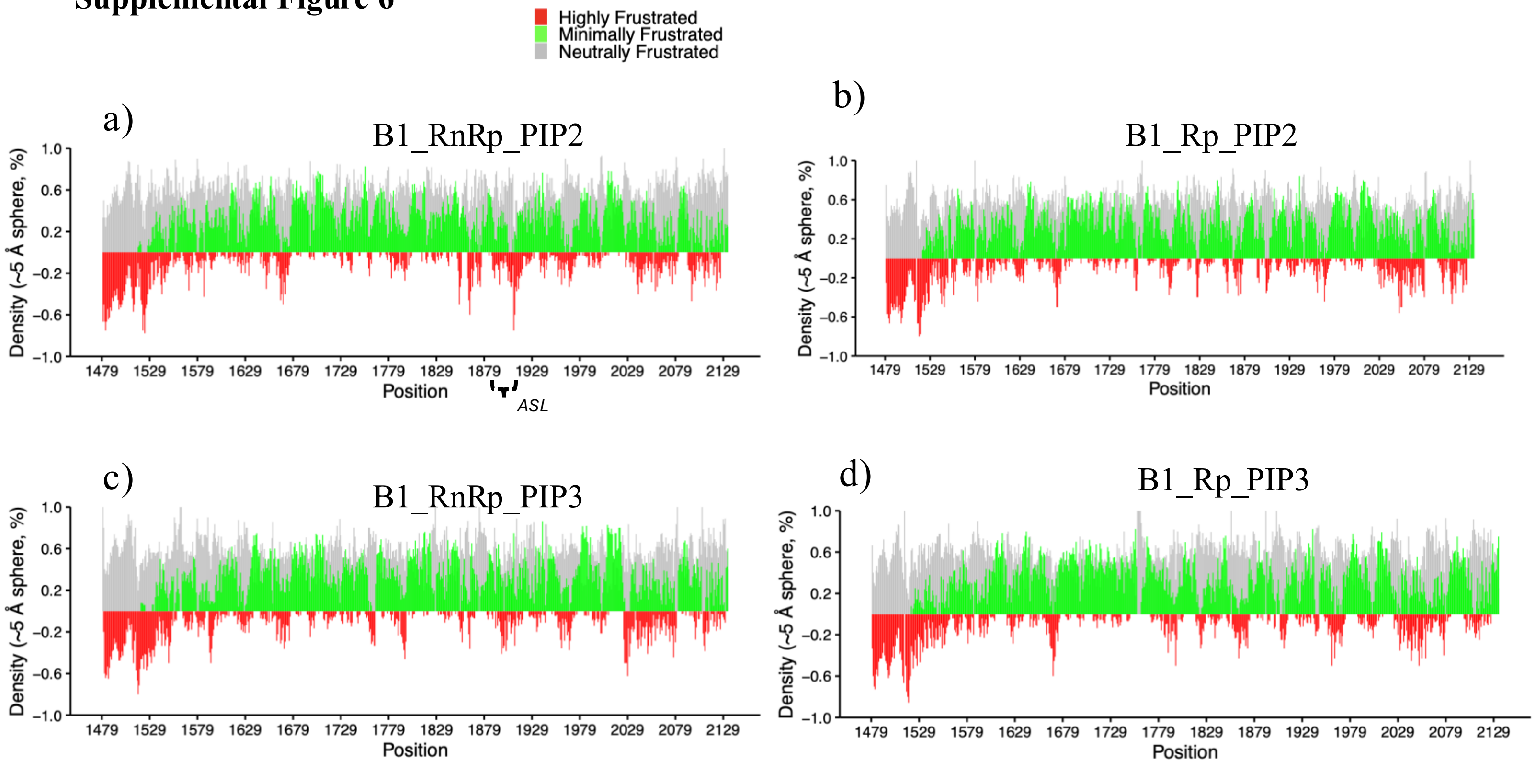


Local frustration analysis of (a) Plexin-B1 in B1_RnRp_PIP2, (b) B1_RnRp_PIP3, (c) B1_Rp_PIP2, and (d) B1_Rp_PIP3.

**Supplemental Figure 7**

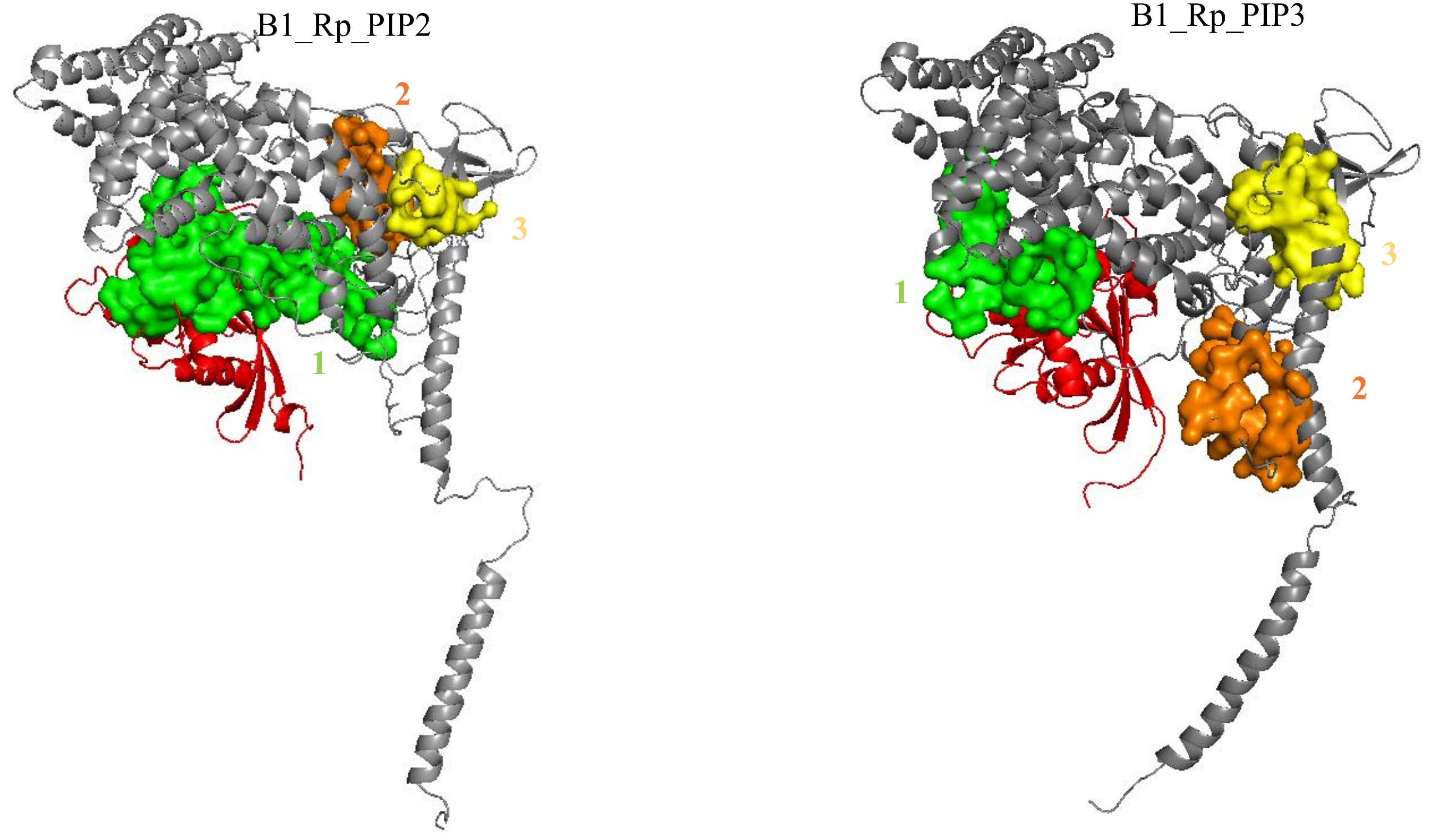


Predicted allosteric site pockets identified on the major cluster structures, with the top three pockets highlighted in distinct colors.
